# Possible planktonic lifestyle of the Silurian trilobite *Deiphon* evaluated through 3D modelling

**DOI:** 10.64898/2026.01.12.698700

**Authors:** Sarah R. Losso, Stephen Pates

## Abstract

Zooplankton are a crucial part of modern oceans, feeding on primary producers and moving nutrients vertically and horizontally. Many euarthropods have evolved planktonic lifestyles, but evaluating extinct taxa were planktonic relies on documenting a broad distribution in diverse lithologies, occurrence across multiple paleobiogeographic regions, and morphological comparisons to modern analogues. Trilobites living in the water column have been proposed to take two forms, well-streamlined and poorly-streamlined, reflecting different lifestyles and behaviors in the water column. Both are believed to go extinct at the end of the Ordovician. A planktonic lifestyle was suggested for *Deiphon* based on its highly inflated, spherical glabella and reduced body, but this is not supported by recent workers. The purpose of the glabellar bubble is also debated, having been both suggested to store low density lipids to increase the buoyancy or host a highly expanded gut. We use a three-dimensional model of *Deiphon* to estimate volume and density of different tissue types (exoskeleton, gut, body, limbs) to calculate its specific gravity. To investigate the impact of the bubble and its contents, we tested the impact on buoyancy by varying presence of the inflated glabella, content of the bubble, and exoskeletal thickness. The sedimentological and geographical information for this animal is poor, making morphology the only proxy available to test its life mode. Exoskeletal thickness has a large impact on the buoyancy of *Deiphon*, suggesting plankton trilobites may have required thin exoskeleton to remain buoyant in the water. Our results support a pelagic lifestyle for *Deiphon*, if its bubble was filled with lipids, which we consider the most likely of the possibilities when the rest of the animal’s morphology is considered.

## INTRODUCTION

Zooplankton are an important part of the modern water column, occupying low trophic levels in marine food webs and forming an integral part of the biological pump (e.g., Turner, 2015; Steinberg and Landry, 2017). In order to remain in the water column, zooplankton employ a range of strategies to maintain their buoyancy, including reducing their density through incorporating light materials such as lipids into their bodies and generating lift through swimming (e.g., Alexander, 1990; Vogel, 1996; Campbell and Dower, 2003). Zooplankton strategies can be divided into several categories: gelatinous floaters; muscle swimmers; and armored hoverers (Tsukamoto et al., 2009). Importantly, these categories include animals that can swim against the prevailing current, thus would not be considered plankton in the strict terms often used by paleontologists (Rigby & Milsom 2000). In this study we treat fossilized pelagic invertebrates as part of the plankton, with the exception of cephalopods, using this broader definition of plankton favored by specialists on extant plankton (e.g., Rigby & Milsom, 2000; Tsukamoto et al., 2009).

Modern plankton use different strategies to remain in the water column. Gelatinous floaters, with a large size range from 1-2 mm in the cnidarian *Muggiaea* to >100 mm in eel larvae), maintain neutral buoyancy with the seawater surrounding them (Tsukamoto et al., 2009). Muscle swimmers, which can also be very small (1-2 mm, such as copepods), but do not reach similar lengths as gelatinous floaters (maximum sizes around 50 mm, such as lobster larvae), have a higher density than their surrounding water, using swimming to generate lift and to maintain their position in the water column (Tsukamoto et al., 2009). Some muscle swimmers are vertical migrants, moving from deeper areas to surface waters to feed during the night (e.g., Bianchi and Mislan, 2016). Armored hoverers generally have the smallest body sizes (from 1 – 25 mm, rarely up to 50 mm), and often have mineralized exoskeletons (e.g., pteropods; (Hunt et al., 2008). These animals have the greatest difference in density with the surrounding water and use swimming appendages to counteract their downward sinking. They sink slowly, due to their small size meaning that they experience seawater as a viscous fluid, and some have large surface areas to provide additional drag to further reduce sinking speeds (Tsukamoto et al., 2009).

Many euarthropod groups have entered the pelagic realm over the Phanerozoic, and this group remains an integral part of the modern zooplankton (Perrier et al., 2015). Inference of a pelagic mode of life in fossil zooplankton requires combining evidence from sedimentology (the animal is found across a range of sediment types), geography (the animal is found across a wide area), and morphology (the animal shares morphological features important for a pelagic mode of life with modern analogues) (e.g., Fortey, 1985). During the Paleozoic examples of euarthropod muscle swimmers include *Isoxys longissimus*, using its carapace shape to generate lift (Pates et al., 2021) and likely large suspension feeding radiodonts such as *Aegirocassis* (Van Roy et al., 2015; Perrier et al., 2015), and examples of euarthropod armored hoverers include trilobite larvae, whose spherical shape and elongate spines are interpreted to have slowed sinking speeds (Laibl et al., 2023). Adult trilobites can be considered as both muscle swimmers and armored hoverers (‘well streamlined’ and ‘poorly streamlined’ groups of Fortey, 1985). ‘Well streamlined’ trilobites are generally larger (up to 100 mm) than the ‘poorly streamlined’ groups (20 – 40 mm), and are interpreted to have generated lift through swimming, while the ‘poorly streamlined’ groups may have utilized elongate genal spines to maintain stability (Fortey 1985) and, by analogy with armored hoverers, slow their rate of sinking.

Pelagic trilobites are considered to be restricted to the late Cambrian and Ordovician (e.g., Fortey, 1985; Adrain et al., 2004). Repeated evolution of a pelagic mode of life in trilobites from over this time period allowed them to take advantage of food sources and habitats, and made them a key link in the early Paleozoic biological pump as zooplankton (Perrier et al., 2015). Computational Fluid Dynamics (CFD) analyses have been conducted on several trilobite species showing a range of lifestyles including the roll of enrollment in swimming (e.g. *Microparia speciosa*; Esteve & López-Pachón 2023), queuing behavior (e.g. *Ampyx priscus*; Trenchard et al., 2017; Song et al., 2021; Wang *et al*. 2024; Trenchard et al., 2025), nektobenthic swimming (e.g. *Hypodicranotus striatus*; Shiino *et al*. 2012, 2014), hopping (e.g. *Placoparia cambriensis*; Esteve *et al*. 2021) and demonstrating sensory function of the anterior glabellae (Gómez-Rodríguez et al., 2025).

While the literature since the 1980s considers adult trilobites living in the water column to be restricted to the late Cambrian and Ordovician (e.g., Fortey 1985; Adrain et al., 2004), some post-Ordovician bubble-headed trilobites, including *Deiphon*, were suggested to have lived in the water column in historic studies from the early 1900s (Fig. 1). The glabellar bubble of this animal was inferred to have been filled with a low-density material (Dollo, 1910; Staff and Reck, 1911; Ruedemann, 1934; Whittard, 1934), such as lipids known from across modern arthropod groups (e.g., Campbell & Dower 2003). Further morphological support comes from the reduced body size relative to the head, suggested as a way to increase the surface area compared to the weight with the development of spines, and thus slow down sinking (Whittard, 1934). Thus, this historically inferred lifestyle is comparable to that of armored hoverers in the modern oceans. *Deiphon* was reinterpreted as a benthic animal following measurement of its exoskeleton thickness at the base of its genal spines (0.30 – 0.35 mm), considered too thick to facilitate low enough buoyancy for a pelagic lifestyle (Holloway and Campbell, 1974). These authors also considered it unlikely that the bubble contained lipids as evidence for this would not be preserved in fossils (Holloway and Campbell, 1974). As the bubble in *Deiphon* is an expansion of the glabella, it is possible this reflects an enlargement of the gut or crop (Fortey and Owens, 1997) although the digestive system is not known in this taxon. There is a broad disparity in the cephalic and thoracic morphology across the polyphyletic grouping of trilobites considered to have ‘bubble-head’ morphologies, and it cannot be assumed that all lived in similar ways (Fortey & Owens 1997). Further progress in interpreting thin section slides and 3D micro-CT scans have allowed us to gain a much better understanding of trilobite morphologies in 3D, and thus allow us to in more accurately estimate their densities (Losso et al., 2023; El Albani et al., 2024; Losso and Ortega-Hernández, 2024). Indeed, trilobite exoskeleton thickness varies significantly across the body, and is often thickest near the doublure (Losso and Ortega-Hernández, 2024), making it likely the exoskeletal thickness of *Deiphon* was much less than 0.3mm for most of its body.

**Figure 1.**
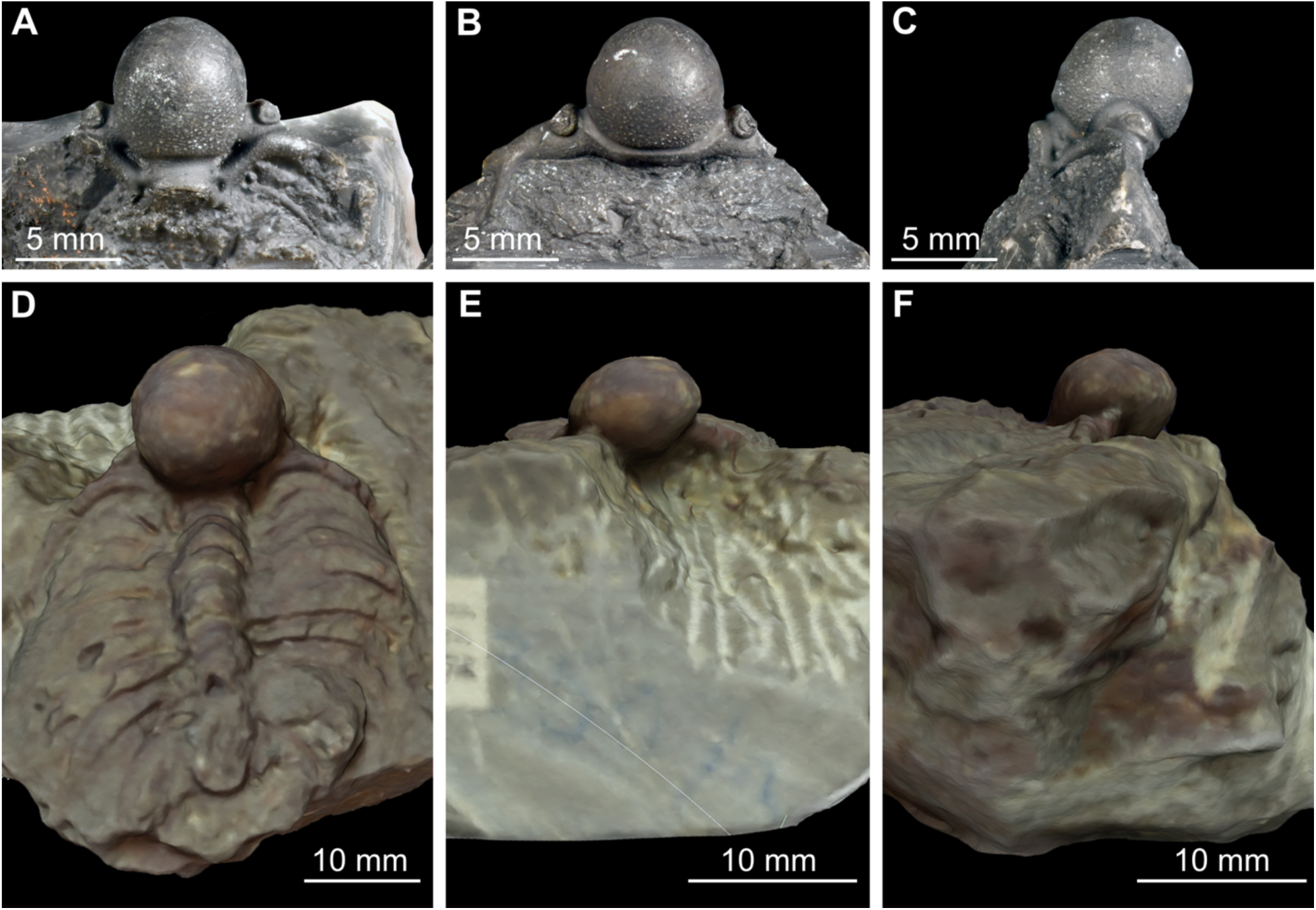
*Deiphon* from the Ordovician of Laurentia and Avalonia with a highly inflated glabella. A – C, USNM PAL 255910 *Deiphon longifrons* Whittard 1934. A, Dorsal view. B, Anterior view. C, Lateral view. D – F, CAMSM A.3507 *Deiphon dikella* Whittard 1934. D, Dorsal view. E, Anterior view. F, Lateral view.

Here, in light of the limited sedimentological and geographical information available for *Deiphon*, we combine volume and density estimates with three-dimensional computational fluid dynamics to evaluate its lifestyle. We consider *Deiphon* lived as an armored hoverer living in the Silurian oceans, providing a robust estimate of its density and considering its stability in the water column.

## METHODS

### Occurrence

*Deiphon* occurrences were downloaded from the Paleobiology Database (PBDB, www.paleobiodb.org) on March 28^th^ 2025 to determine the range of lithologies from which *Deiphon* has been recovered. Lithology data from “lithology 1” and “lithology 2”, and depositional environmental data from the “environment” was simplified following Bault *et al*. (2022), in order to link lithologies to water depth and distance from shore.

### Model creation

A 3D model of *Deiphon dikella* specimen CAMSM A.3507 housed at the Sedgwick Museum of Earth Sciences (Cambridge, UK; CAMSM) was constructed from photographs using photogrammetry in Agisoft Metashape 1 (www.agisoft.com). A three-dimensional model of the exoskeleton of *Deiphon dikella* was created based on CAMSM A.3507 using Shapr3D (https://www.shapr3d.com/). To test the impact of the glabella bubble on the buoyancy of *Deiphon*, two models were created, one with a highly inflated bubble and the other with a glabella similar to *Ceraurus pleurexanthemus* Green 1832. Non biomineralized tissues based on closely related other cheirurid trilobites housed at the Museum of Comparative Zoology at Harvard University (Cambridge, USA; MCZ) including *Ceraruus plaurexanthemus* (MCZ:IP:110933 and MCZ:IP:112018) and *Anacheirurus adserai* (MCZ:IP:202508) were added to both models, also using Shapr3D.

### Volume estimates

The surface area of the exoskeleton without the inflated glabella was estimated by finding the area of librigena, genal spine, half of the thorax and pygidium in ImageJ (Schneider et al., 2012) and multiplying by two. *Deiphon* has an originally poorly vaulted exoskeleton (Fig. 1D – F). The exoskeletal thickness of *Deiphon* is unknown for the majority of the body, and so three thicknesses were used in order to explore the importance of this feature for the overall density of *Deiphon*.

Four values of exoskeleton thickness were used to capture the possible range, 0.008 mm, 0.100 mm, 0.175 mm, and 0.300 mm, drawn from comparisons with closely related cheirurid *Ceraurus pleurexanthemus* and trilobites from the Ordovician of Spitsbergen, which show a broad range in exoskeletal thicknesses across Trilobita (Fortey and Wilmot, 1991; Losso and Ortega-Hernández, 2024). The volume of the exoskeleton without the inflated glabella was calculated by multiplying the area by each potential thickness (Fig. 2; Supplemental Table 1). The inflated glabella exoskeleton volume was found by creating internal spheres by subtracting the exoskeleton thickness from the measured size (Fig. 2; Supplemental Table 1). The volume of the inner sphere of the glabella was calculated by removing the exoskeleton thickness from the radius of the larger sphere (Fig. 2).

**Figure 2.**
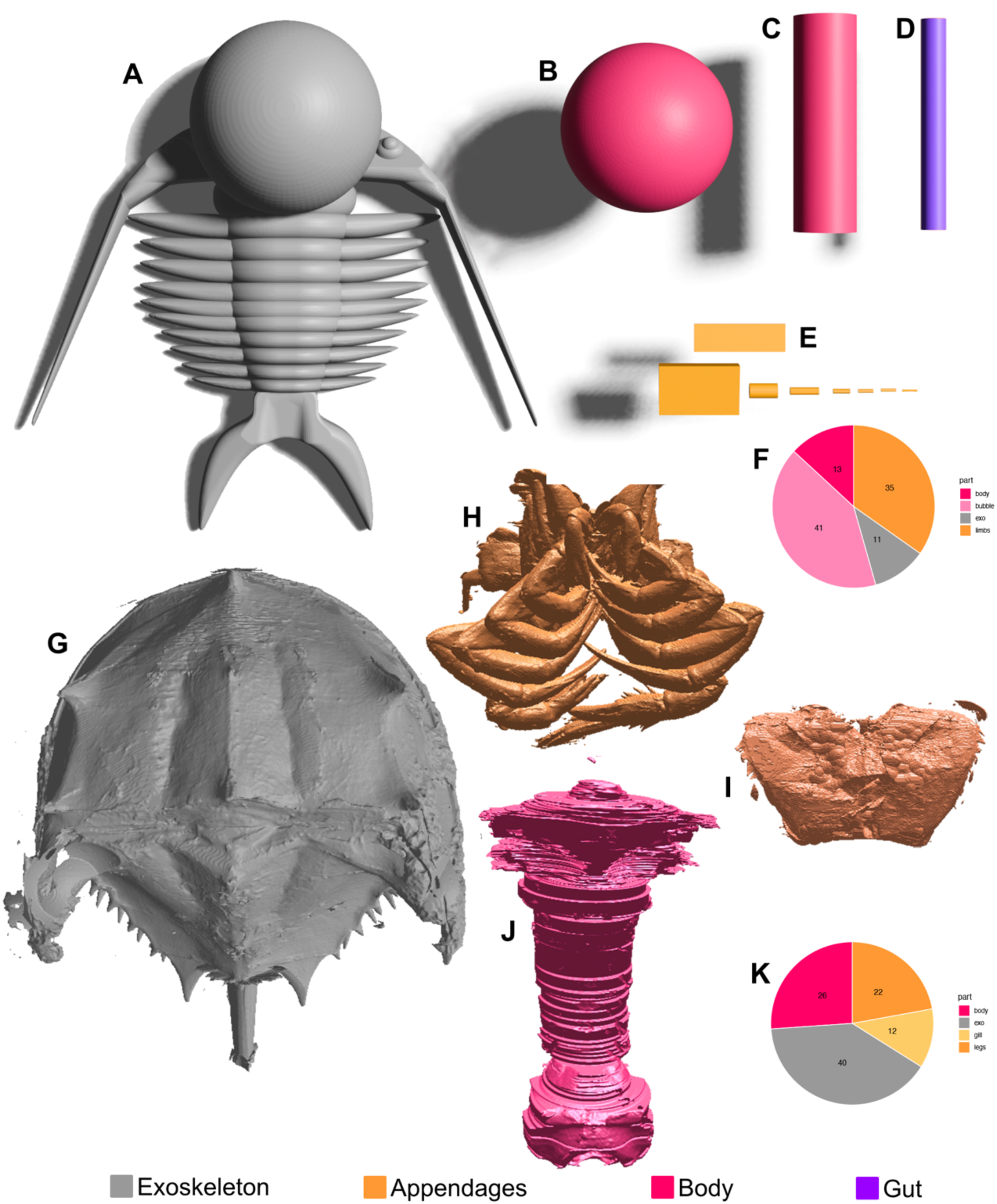
Volume estimates of exoskeleton and non-biomineralized tissues in trilobites and horseshoe crabs. A – E, Volume estimates of *Deiphon dikella* based on A3507. A, Model of exoskeleton morphology. B, Internal volume of glabella bubble. C, Volume of body estimated as a cylinder with a diameter the width of the axial lobe. D, Volume of the gut tract estimated as cylinder occupying 20% of the axial lobe width. E, Volume estimate of biramous appendage with wedge shaped protopodite, cylindrical podomeres and cuboid exopodite. F - I, 3D reconstruction of *Limulus polyphemus* MCZ:IP:41112. F, Dorsal view of exoskeleton. G, Ventral view of cephalothoracic appendages. H, Ventral view of cephalothoracic appendages. I, Ventral view of body lumen.

The volume of non-biomineralized tissues was based on comparison with *Ceraurus pleurexanthemus* where the body cavity occupies a cylindrical space under the axial lobe which is as deep as it is wide (Losso and Ortega-Hernández, 2024). A thin membrane extends from the body cavity to the doublure, but is only as thick as the exoskeleton, so it was not included in our volume estimates. The axial lobe of A3507 is ca. 2.5 mm across and only slightly tapers posteriorly along the ca. 10 mm length, so a cylinder with a radius 1.25 mm was used to estimate the volume of the body (Fig. 2). The gut was estimated to occupy 20% of the width of the axial lobe based on the cheirurid *Anacheirurus adserai* MCZ:IP:202508 from the Fezouata Shale. The gut was modeled as a cylinder with a radius of 0.5 mm and length of 10 mm (Fig. 2; Supplemental Table 1).

The volume of appendages were estimated based on the two cheirurids with known preserved limb morphology, *Anacheirurus adserai* and *Ceraurus pleurexanthemus* (Pérez-Peris et al., 2021; Losso and Ortega-Hernández, 2024). Podomere numbering follows recent studies of extant arthropods showing a unified framework (Bruce and Patel, 2018, 2020; Bruce, 2021). The protopodite is wedge shaped in cross section being sagittally broadest dorsally and tapering ventrally (Losso et al., 2023; Losso and Ortega-Hernández, 2024). A rectangular wedge was used to estimate the volume of the protopodite, 1.14 mm width 1.99 mm in height, and 3.19 mm length (Fig. 2; Supplemental Table 1). Podomere dimensions vary between trilobite species (Pérez-Peris et al., 2021; Losso and Ortega-Hernández, 2022, 2024), but can be estimated as cylinders with decreasing diameters distally (Fig. 2). The proximal three podomeres were based on measurements from *C. pleurexanthemus* (MCZ:IP:110933 and MCZ:IP:112018; Losso and Ortega-Hernández, 2024) whereas the distal podomeres were determined by the proportions see in *A. adserai* (Pérez-Peris et al., 2021; Supplemental Table 1). The exopodite was estimated as a cuboid the size of the exopodite articles since the volume of the lamellae is miniscule. The length (3.62 mm) and width (1.16 mm) of the cuboid were based on *A. adserai* (Pérez-Peris et al., 2021) and the height (0.13 mm on *C. pleurexanthemus* Losso and Ortega-Hernández, 2024; Supplemental Table 1). Trilobite appendages decrease in size along the axis, with the posteriormost appendages of *Olenoides serratus* being 60% the volume of the anteriormost (Supplemental Table 1).

### Density estimates

In order to estimate the density of *Deiphon,* the density of the individual body parts (exoskeleton, interior of bubble, and soft anatomy) was required. For the exoskeleton, the density of calcite was used (2.711 g/cm3) (DeFoe and Compton, 1925). For the interior of the bubble, both lipids (0.00091 g/cm3) (Lee and Joye, 2006)and sea water (1.03 g/cm3) were used. For the gut, the density of sediment (0.8 g/cm3) was used.

For the soft parts, comparisons to the density of the soft parts of modern crustaceans were used. These were calculated using densities of three types of material that make up the composition of arthropod soft tissues: water (1 g/cm3), lipids (0.92 g/cm3), and proteins (1.45 g/cm3) (values from Moya-Laraño et al., 2008). The mass of water, protein (calculated as the nitrogen content multiplied by 6.25), lipid and ash (density 0.61 g/cm3) for 13 shrimps, lobsters, and prawns were collected from the USDA website. The known density of each tissue or ash was then used to calculate the overall density of the soft tissues of each of these samples. The mean density across all 13 samples (1.026 g/cm3) was used as the density for the soft parts of *Deiphon* (Supplementary Excel Spreadsheet 2).

The estimated volumes for each body part (exoskeleton, bubble interior, gut, and soft parts) were multiplied by their estimated densities and divided by the total volume to give an overall density for the *Deiphon* model. The specific gravity was calculated using equation 1, for comparison to extant pelagic and planktonic animals. A density of seawater of 1.03 g/cm3 was used (www.engineeringtoolbox.com).

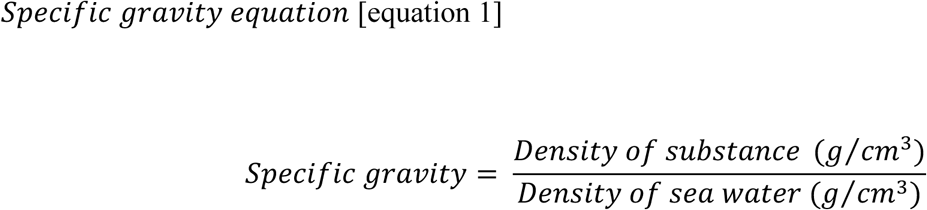

### Computational fluid dynamics

A three-dimensional model of *Deiphon* based on A3507 was made in Shapr3D and exported as .stl type. Mesh creation, geometry setup, and fluid simulations were performed using ANSYS Workbench and DesignModeler, Meshing, and Fluent components, all version 2024 R2.

Models were imported into the ANSYS SpaceClaim environment, shrinkwrapped, and converted to solid objects of the proprietary ANSYS filetype “.scdoc”. SpaceClaim models were then imported into the ANSYS Workbench DesignModeler, and scaled to 0.02 m long, comparable to the length of fossil specimens (e.g. Fig. 1). A virtual flume tank of dimensions 0.05 x 0.05 x 0.4 m was constructed using sketch planes which were then converted to a solid and then extruded in the Z dimension to create a three-dimensional cuboid. These dimensions were chosen to be large enough so that the fluid flow would not interact with the margins of the tank, as the model was positioned 0.025 m from the sides of the tank, 0.1 m from the inlet and 0.3 m from the outlet. Sketch planes and the extrude function were also used to create a smaller cuboid 0.02 x 0.02 x 0.4 m inside the tank for the creation of a volume with smaller mesh elements during the meshing step.

A series of meshing controls were applied to create a coarse, medium, and fine mesh, with the number of elements ranging from ∼750,000 to ∼3.1 million (Table 1). An Element size control was applied to the whole mesh to set the maximum size for a mesh element. The size varied across the coarse, medium, or fine mesh (Table 1). A Patch Conforming Method was applied to the whole geometry, to create a tetrahedral shape for mesh elements. A Body Sizing was applied to the smaller cuboid around the *Deiphon* model, extending to the anterior and posterior of the virtual flume tank, to set the maximum size for a mesh element close to the model and in its wake. Again, a range of sizes were applied, for the coarse, medium, and fine meshes respectively (Table 1). Face sizing was applied to the surface of the *Deiphon* model, to capture the details of the model by setting a minimum face size for elements touching the model. Three different sizes were used, one each for coarse, medium, and fine meshes (Table 1). An Inflation layer was also applied to the surface of *Deiphon* model, to create small elements that capture the boundary layer close to the model. 20 layers were used with the ‘smooth transition’ inflation option, with a transition ratio of 0.272, and growth rate of 1.2.

**Table 1.** Mesh sizes in CFD simulations.

|  | Coarse | Medium | Fine |
| --- | --- | --- | --- |
| Element size | 2 e-2 m | 1 e-2 m | 9e-3 |
| Body sizing | 5.5 e-3 m | 2.5e-3 m | 1.5e-3m |
| Face sizing | 1.25 e-3 m | 5e-4 m | 4e-4m |
| Total number of elements | 749795 | 1361016 | 3092750 |

Fluid flow simulations were performed using ANSYS Fluent. The pressure-based solver with absolute velocity formation and steady state were used, with the viscous realizable k-epsilon turbulent solver, with enhanced wall treatment. The k-epsilon model is a two equation model, based on transport equations for the turbulent kinetic energy (k) and dissipation rate (epsilon) (Launder and Spalding 1972). The realizable k-epsilon model differs from the original by modifying the transport equation for the dissipation rate and providing an alternative formula for the turbulent viscosity (Shih et al., 1995). Further details on the mathematics behind the models and their implementation in ANSYS is provided in the documentation (ANSYS Academic Fluent, release 2024 R2)

The fluid was set to have the properties of water-liquid in the Fluent Database (density 998.2 kg/m^3^; viscosity 0.001003 kg/ms. The inlet velocity was set to 0.1m/s, simulating a slow swimming speed for *Deiphon,* and the pressure outlet was set to zero gauge pressure.

Simulations were run for all three mesh types, to ensure that interpretations were mesh independent. At least 100 iterations were run, and more iterations were run and until the solution was converged. Simulations were treated as converged if residuals of all six equations (continuity, x-velocity, y-velocity, z-velocity, k and epsilon) were all below 1 × 10^−6^. Although not interpreted, values for drag and lift force acting on the model were also collected, to confirm that these were also stable and unchanged across at least 50 iterations, for a simulation deemed converged.

The wall Y+, contours of pressure, and vectors of velocity were visualized for the surface of the *Deiphon* model (wall Y+) and along the Z axis through the midline of the *Deiphon* model (pressure, velocity) to ensure the Y+ was low enough for the k-epsilon model to provide reasonable results for the boundary layer, and to visualize the impact of the *Deiphon* model on the flow of water around the model.

## RESULTS

### Occurrence

Specimens assigned to the genus *Deiphon* have been recovered from a broad range of environments, including shallow marine, reefal, slope/basin, and the outer platform (Fig. 5A). There are slightly more occurrences of the genus in deeper water environments than shallow ones (Fig. 5A). Sixteen of the occurrences on PBDB are only identified as marine. *Deiphon* specimens are found in nine different lithologies (Fig. 5B). The most common lithology is limestone, followed by shale. There are five occurrences of *Deiphon* in each dolomite, lime mudstone, and grainstone, three each in marl and sandstone, and two each within mudstone and siltstone (Fig. 5B). *Deiphon globifrons* is known from shallow and outer platform environments while other species are restricted to a single environment (Supplemental Table 3).

### Buoyancy in Deiphon

The specific gravity of *Deiphon* is largely controlled by the estimated exoskeleton thickness, material within the glabellar bubble and presence/absence of the bubble (Fig. 3). Filling the bubble with lipids results in a 0.06-0.08 specific gravity decrease across the different exoskeleton thickness treatment groups. The thickest exoskeleton estimate, 0.3mm, results in specific gravity ranging from 1.385 – 1.446 with sea water resulting in the highest value (Supplemental table; Fig. 3). The medium exoskeleton (0.175mm) results in a more buoyant body with specific gravity ranging from 1.217 – 1.287 (Supplemental table; Fig. 3). The exoskeleton of 0.1 mm based on *Ceraurus pleurexanthemus* results in a body with specific gravity ranging from 1.094 – 1.142 (Supplemental table; Fig. 3). The thinnest exoskeleton (0.008mm) leads to the most buoyant body, specific gravity 0.954 – 1.040, with lipid the lipid filled bubble being more buoyant than surface sea water at 10°C (Supplemental table; Fig. 3). Within the thick and medium exoskeleton treatment groups, the absence of a glabellar bubble results in higher specific gravity (Supplemental table; Fig. 3). The impacted of the bubble is more nuanced with the thinnest exoskeleton. The sea water filled glabella results in similar specific gravity with or without a glabellar bubble (Supplemental table; Fig. 3). But with lipid filled, the bubble significantly decreased specific gravity (Fig. 3). The thin exoskeleton results in nearly neutrally buoyant bodies (sea water filled bubble, and the two no bubble). The lipid filled bubble with a thin exoskeleton has positive buoyancy compared to sea water.

**Figure 3.**
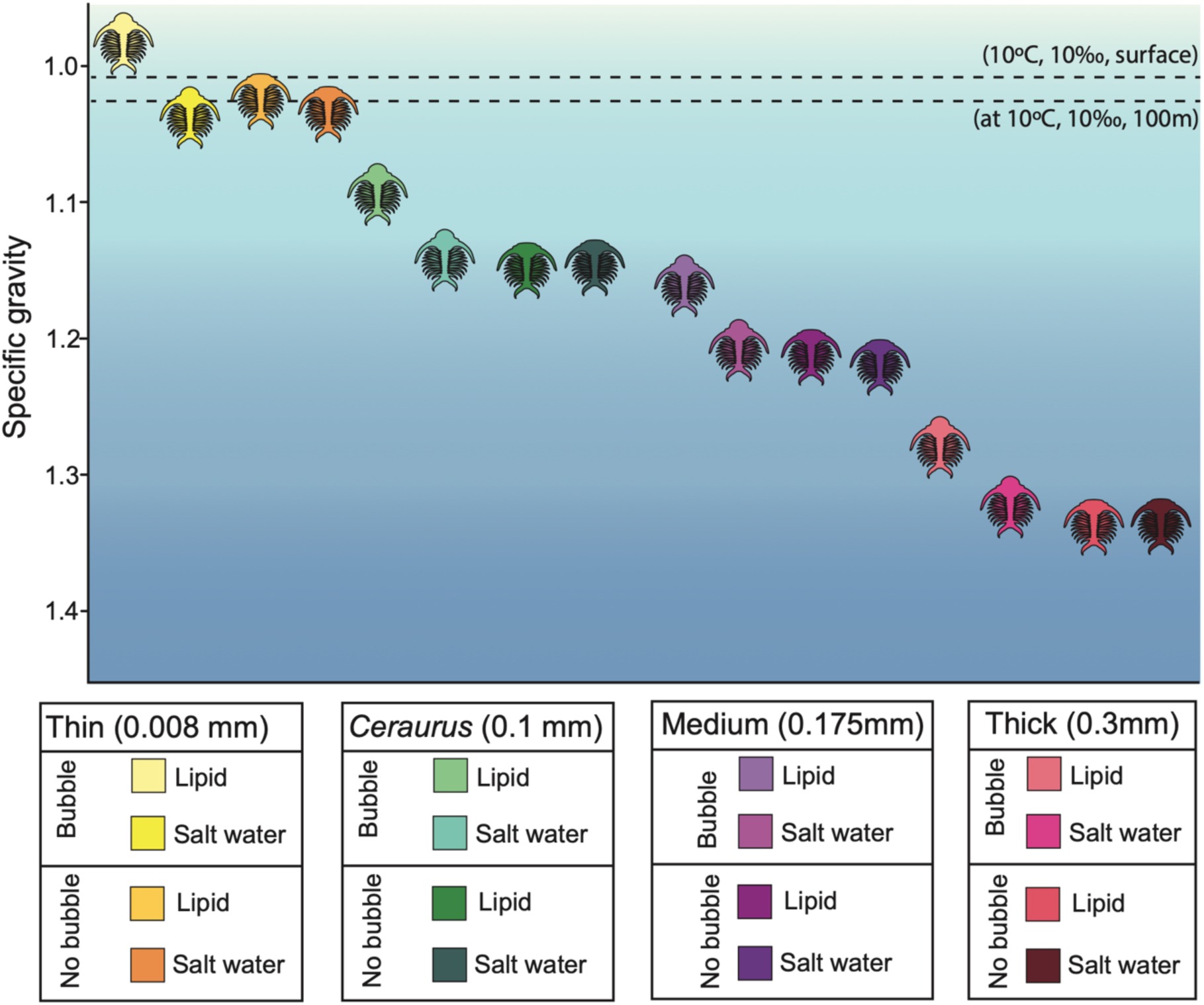
Buoyancy of *Deiphon* with bubble-shaped glabella and without.

**Figure 4.**
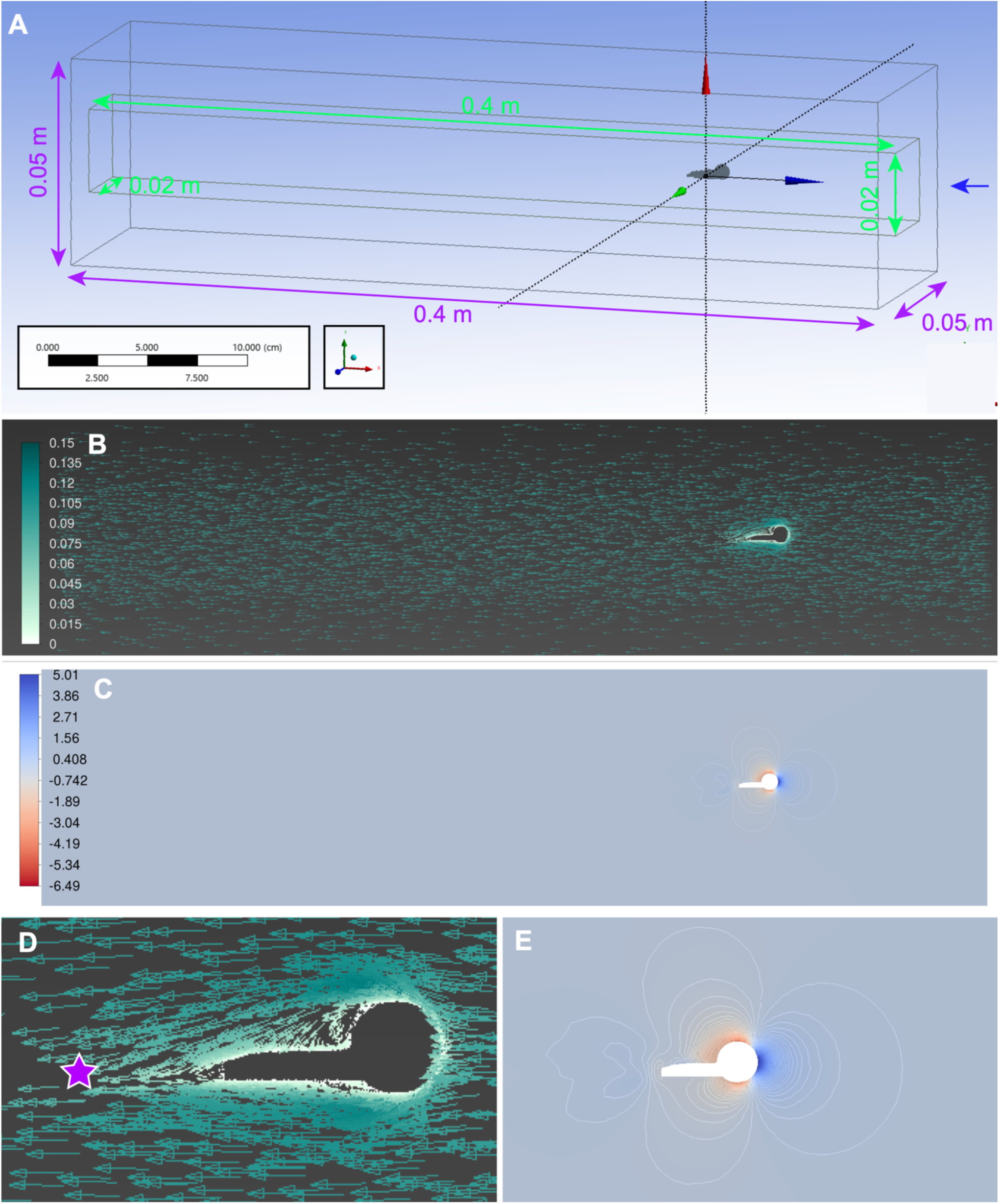
CFD simulation results of prone *Deiphon* in water column. **A,** Geometry of virtual flume tank with dimensions, body of influence, and position of model. Blue arrows show inlet. **B,** Velocity field for CFD simulation using medium mesh (m/s). **C,** Pressure contours for CFD simulation using medium mesh (Pa). **D,** Magnification of velocity field around body (m/s). **E,** Magnification of pressure contours around body (Pa).

**Figure 5.**
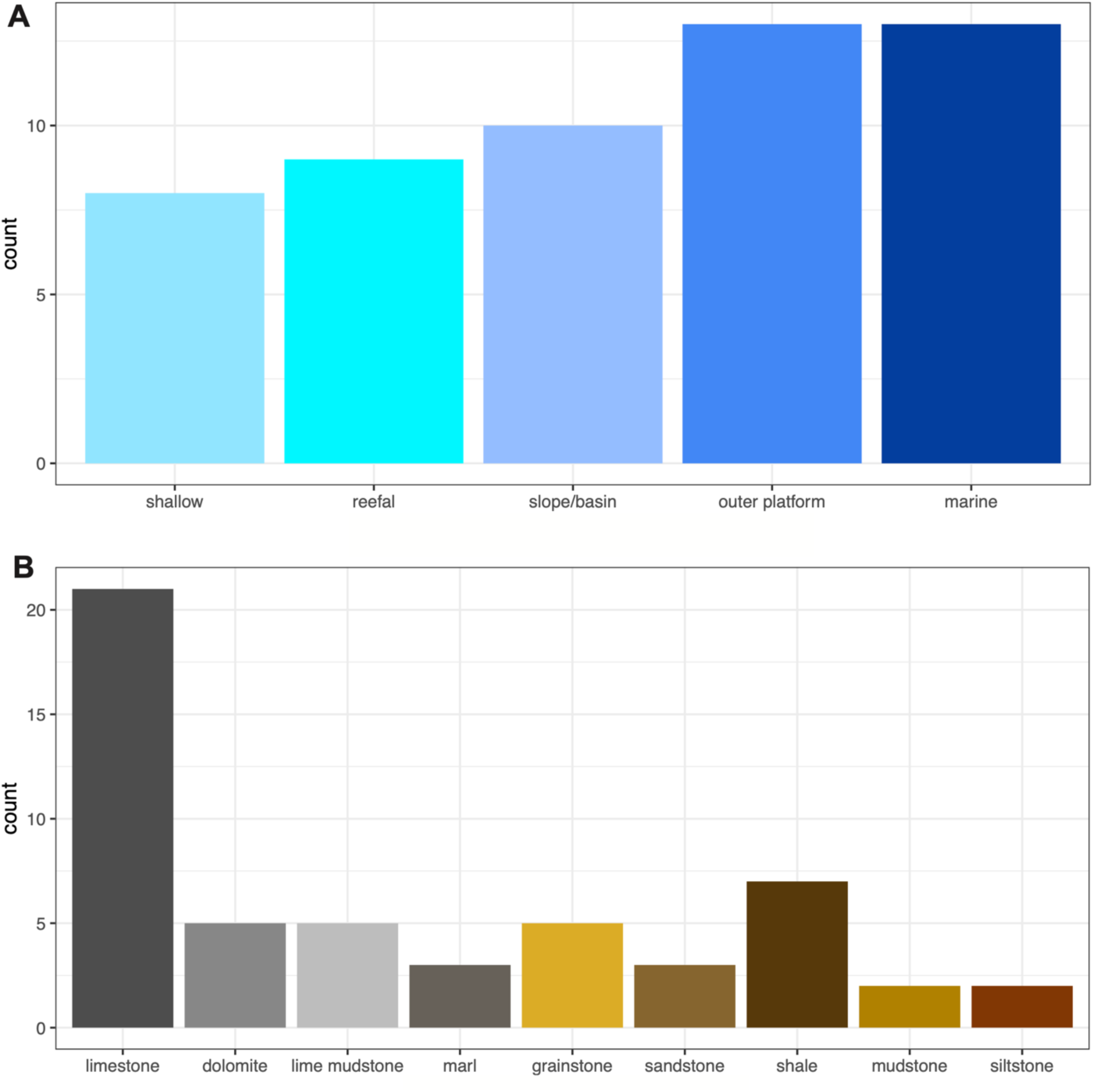
Occurrence of *Deiphon*. **A,** Occurrence of *Deiphon* by depositional environment. **B,** Occurrence of *Deiphon* by lithology.

The thick and medium exoskeletons without a glabellar bubble result in specific gravities that are lower than those of modern plankton (Fig. 6). The groups with the two thinnest exoskeletons and glabellar bubble filled with lipids all fall within or near specific gravity range of modern plankton (Fig. 6). Thick and medium exoskeletons with a glabellar bubble fall within the range of armored hoverers, but those organisms are much smaller (Fig. 6).

**Figure 6.**
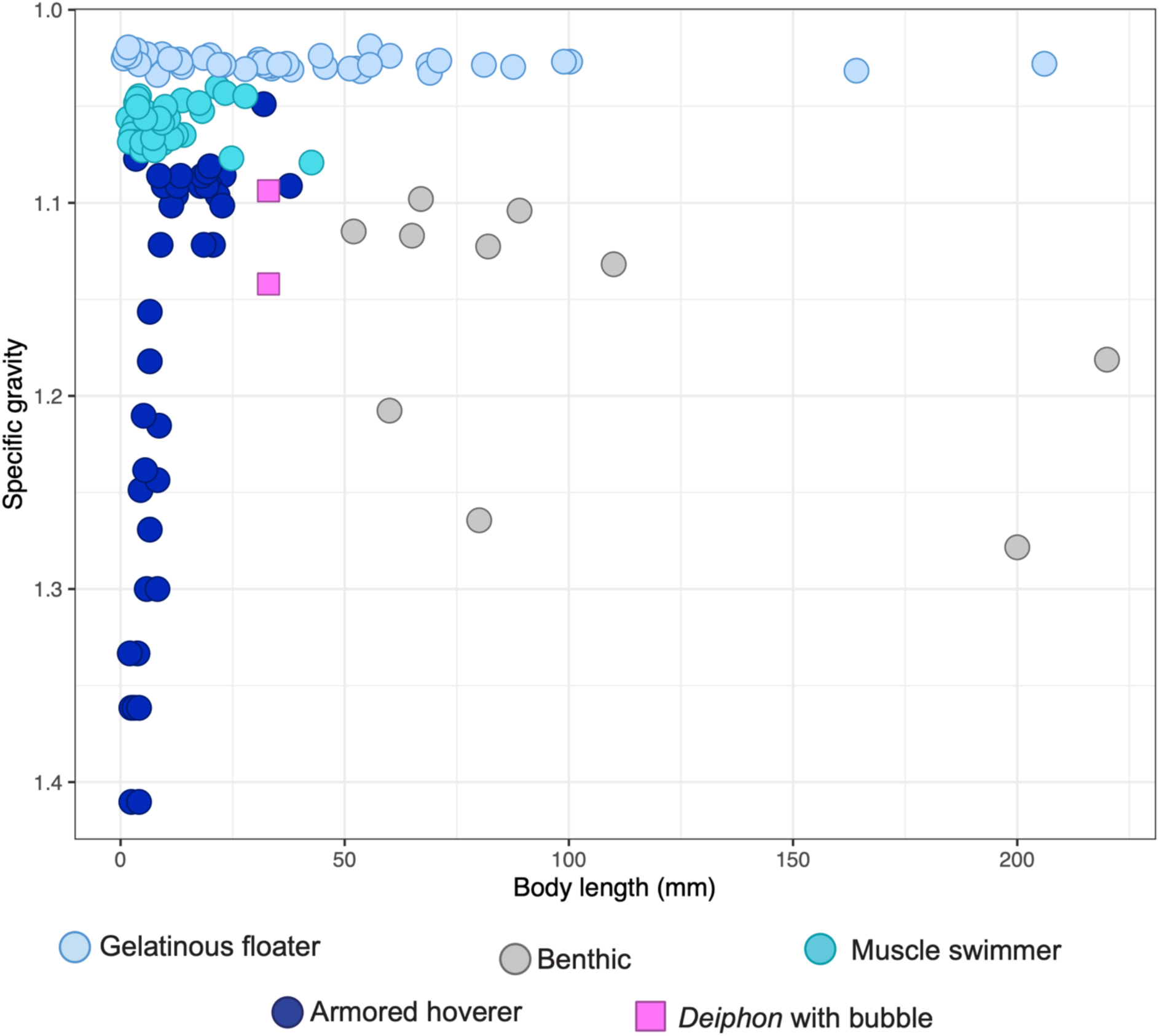
Specific gravity compared to body size of planktonic metazoans. Planktonic data was digitized from (Tsukamoto et al., 2009) and benthic data was added from (Spaargaren, 1979).

### Stability in fluid flow

In all cases there was no interaction between diverted fluid flow and the edges of the fluid domain, meaning that the domain was large enough to avoid edge effects. Wall Y+ values for all three mesh types were <1 across the surface of the *Deiphon* model (Supplementary Figure 2). Results were consistent across the three mesh types (coarse, medium, fine), and so interpretations can be considered mesh and domain size independent (Supplementary Figures 3).

The medium mesh type is visualized in Figure 4 (coarse and fine meshes in Supplementary Figure 3). Flow is initially uninterrupted, but the fluid velocity slows as it encounters the anterior margin of *Deiphon* (Fig. 4A, B). Fluid flow remains slow near the surface of the model, particularly posterior to the glabellar bubble. The field velocities show a small vortex immediately behind the glabella and a wake less than the length of body, with flow recombining ∼ 15 mm behind the posteriormost point of the *Deiphon* model (Fig. 4D). No vortexes form below or behind the body.

A similar profile can be seen when visualizing the pressure contours. An area of high-pressure forms in front of the body, at the tall anterior edge of the glabellar bubble (Fig. 4C, E). A low-pressure region forms dorsal to the glabellar and body. This ranges from near the dorsal margin of the bubble and extends over the first third of the thorax (Fig. 4E). Another smaller low-pressure region forms in a similar position ventral to the cephalon but does not extend below the thorax (Fig. 4E).

## DISCUSSION

### Estimating the density of trilobites: a new approach

The density of a body will depend on the constituent parts, both the materials of each and the proportions within the body. Previously, the density of modern marine benthic crustaceans has been used for trilobites (1.14g/cm^3^; Wang et al., 2024), which assume a homogenous body. Trilobite are composed of several different materials, the high-density calcite exoskeleton (0.0027 g/cm^3^), the body (0.001 g/cm^3^), and the gut which could be filled with various materials (Supplemental Table 2). Event small increases in exoskeletal calcite thickness have dramatic impacts on density.

The density of *Deiphon* is controlled by exoskeletal thickness, resulting in specific gravities (SG) ranging from less than sea water to similar as planktonic animals (Figs. 3, 6). The impacts on density caused by increases in exoskeletal thickness would force tradeoffs between being heavily armored for defense or light weight for planktonic lifestyles. The bubble dramatically changes the volume of the body, occupying over half of volume of the individual (Supplemental Figure 1), thus the presence and material within the bubble both impact SG. Only the thinnest exoskeleton treatment of *Deiphon* results in a neutrally buoyant body, suggesting this would rarely occur in other trilobites lacking specialized morphology. The exoskeleton thickness at the base of *Deiphon’*s genal spine was reported as 0.30 – 0.35 mm (Holloway and Campbell, 1974), much thicker than the range (0.07 – 0.13 mm) reported from Spitsburgen trilobites across a range of environments (Fortey and Wilmot (1991). However, the exoskeletal thickness has not been investigated over the rest of the body of *Deiphon*. If the thickness of 0.3 mm was maintained across the entire body, it would significantly weigh down the body indicating a benthic lifestyle (Fig. 3). However, a thickness of 0.3 mm is unlikely, as the exoskeletal thickness was likely thicker at the base of the genal spine than across most of the exoskeleton. The closely related *Ceraurus pleurexanthemus* shows a much thicker exoskeleton (0.3 mm) close to the doublure, where it interacts with sediment, than for the rest of the body (0.1 mm). Thus, it is reasonable to consider *Deiphon* with a similar exoskeletal thickness to *C. pleurexanthemus* at 0.1 mm.

Non-biomineralized tissues are not known in *Deiphon*, thus there is no directed evidence for material occupying the glabellar bubble. There are two competing theories for material filling the bubble, either the expanded gut Fortey and Owens, (1997) or a low-density material such as lipids (Dollo, 1910; Staff and Reck, 1911; Ruedemann, 1934; Whittard, 1934).

An expansion of the gut in *Deiphon* to occupy the entire glabella would likely reflect a highly enlarged crop, which in modern crustaceans is used to masticate food (Ceccaldi, 1989; Rupert and Barnes, 1994; Lerosey-Aubril et al., 2011; Hopkins et al., 2017) (Ceccaldi 1989; Ruppert & Barnes 1994). Although crops have evolved in trilobites multiple times, convincing evidence for them is not wide spread among different taxa (Lerosey-Aubril et al., 2011). Although a crop were reported in the cheirurid *Ceraurus pleurexanthemus,* this has since been regarded as unlikely to represent a preserved digestive system (Lerosey-Aubril et al., 2011). The functional implications of such an enlarged crop are unclear, as it may make mastication difficult. The ventral side of the glabellar bubble in *Deiphon* is unknown, but evidence of muscle attachment would support the presence of a crop (Fortey and Owens, 1997; Lerosey-Aubril et al., 2011).

Modern plankton often store lipids within the body both as energy storage and to regulate buoyancy (Lewis, 1970; Lee and Hirota, 1973; Childress and Nygaard, 1974; Sargent and Falk-Petersen, 1988). Unlike in modern copepods where the lipids are easily compressed at depth (Campbell and Dower, 2003), the rigid calcite exoskeleton of *Deiphon* would prevent this. The storage of lipids could have aided *Deiphon* in surviving long periods without food or dispersal and to increase buoyancy. We regard lipid storage as the more likely function of the glabellar bubble as a crop of this size would struggle to masticate food since the lumen would be very large.

### Planktonic lifestyle for Deiphon

Fortey (1985) suggested three criteria to infer a planktonic lifestyle for an extinct organism: 1) occurring in a broad range of sediment types, 2) occurring in a wide geographic area and 3) morphology resembling modern analogs. The limited number of specimens known for *Deiphon* with only sixteen occurrences on PBDB, make it challenging to use sedimentological and paleobiogeographical to assess the planktonic lifestyle. Planktonic animals are expected to occur in a broad range of environments because their lifestyles facilitate dispersal, but with limited numbers of individual specimens, it is unlikely for them to be found in many locations. Deiphonines are distributed broadly through the low to intermediate latitudes (Pérez-Peris et al., 2024). *Deiphon* is known from a wide range of environments and lithologies, suggesting a pelagic lifestyle that facilitated dispersal (Fortey, 1985). However, only one species, *Deiphon globifrons* is known from shallow and outer platform environments while other species all occur within one formation, with very limited sample sizes (Supplemental Table 3). Considering this limited sedimentological and paleobiogeographic information available for *Deiphon* since it is not a highly abundant taxon, the morphology is the strongest evidence to evaluate its lifestyle.

The buoyancy of adult trilobites has not previously been estimated. Morphological features hypothesized to increase buoyancy across trilobites and closely related groups include elongate spines and a highly vaulted exoskeleton in larval trilobites (Speyer and Chatterton, 1989; Park and Choi, 2010; Laibl et al., 2023), and the thin exoskeleton of the nektaspid *Naraoia* (Haug and Haug, 2016). The primary control on trilobite buoyancy comes from the exoskeletal thickness (Figure 3).In addition, the lower SG for *Deiphon* models with bubbles compared to those without, for all exoskeleton thicknesses considered, demonstrates that the presence of the highly inflated glabella increases the buoyancy of *Deiphon* regardless of the infilling fluid, providing a secondary control on buoyancy for bubble-headed trilobites.

For trilobites traditionally considered planktonic, the exoskeletal thicknesses likely varied depending on if they were muscle swimmers or armored hoverers, as these require different buoyancies. The well streamlined muscle swimmers require lower SG than armored hoverers, and thus are expected to have thinner exoskeletons leading to lower overall densities. *Deiphon*’s estimated specific gravity of ∼1.05, combined with its reduced body with large spines support an inference of the armored hovered planktonic mode. Modern animals such as *Gigantocypris*, a modern deep see giant copepod, has a nearly spherical body possible analogous to the large bubble-head of *Deiphon*, and swims slowly by ‘dragging’ itself through water at low Reynolds numbers, using its second antennae (Davenport, 1990). We speculate that *Deiphon* may have used its limbs sparingly to provide thrust and lift, counteracting its slow sinking. The crucial role of exoskeletal thickness on potential lifestyle highlights the need for further study in more clades and how it changes across the body. This possible reinvasion of trilobites into the water column after the End Ordovician Mass Extinction, evidenced by *Deiphon*, increases the range of known lifestyles and ecological roles for members of this diverse arthropod clade into the Silurian. Importantly, considering the range of different morphologies and planktonic strategies available in modern oceans and this model-based approach to estimate the densities of trilobites may facilitate recognition of additional trilobite armored hoverers during other periods in the Paleozoic.

## Acknowledgements

We thank Matt Riley (Sedgwick Museum, Cambridge) for access to the collections and loan of camera equipment for photography of *Deiphon dikelus*. We thank Pauline Affatato (Columbia University) for providing the CT scan of *Limulus polyphemus*, MCZ:IP:41112; Gene Hunt (Smithsonian Institution), and Conrad C. Labandeira (Smithsonian Institution) for facilitating access to specimens. SP acknowledges funding from a NERC Independent Research Fellowship NE/X017745/2.

## Supplemental Material

**Supplemental Figure 1.**
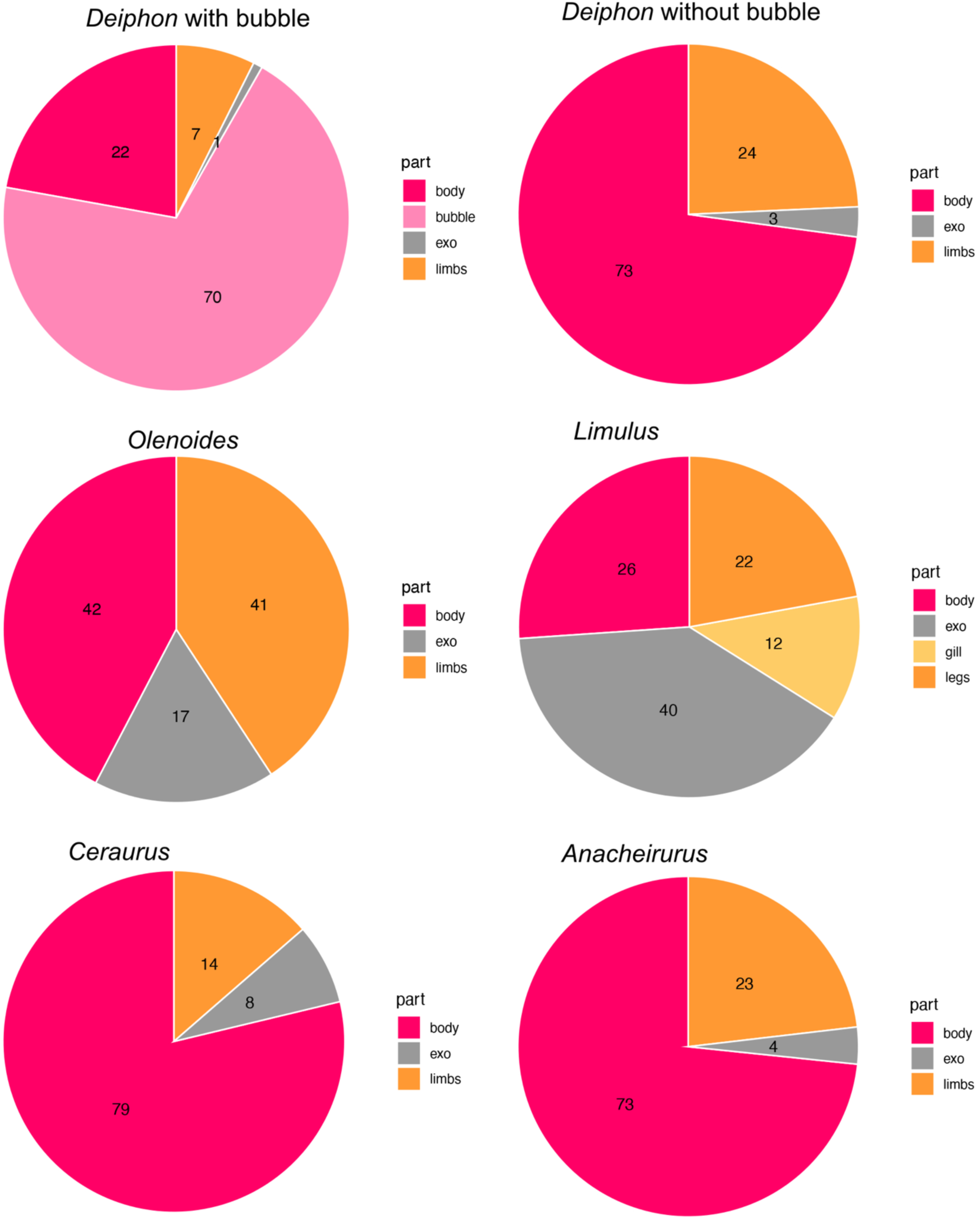
Body volume proportions in euarthropods.

**Supplemental Figure 2.**
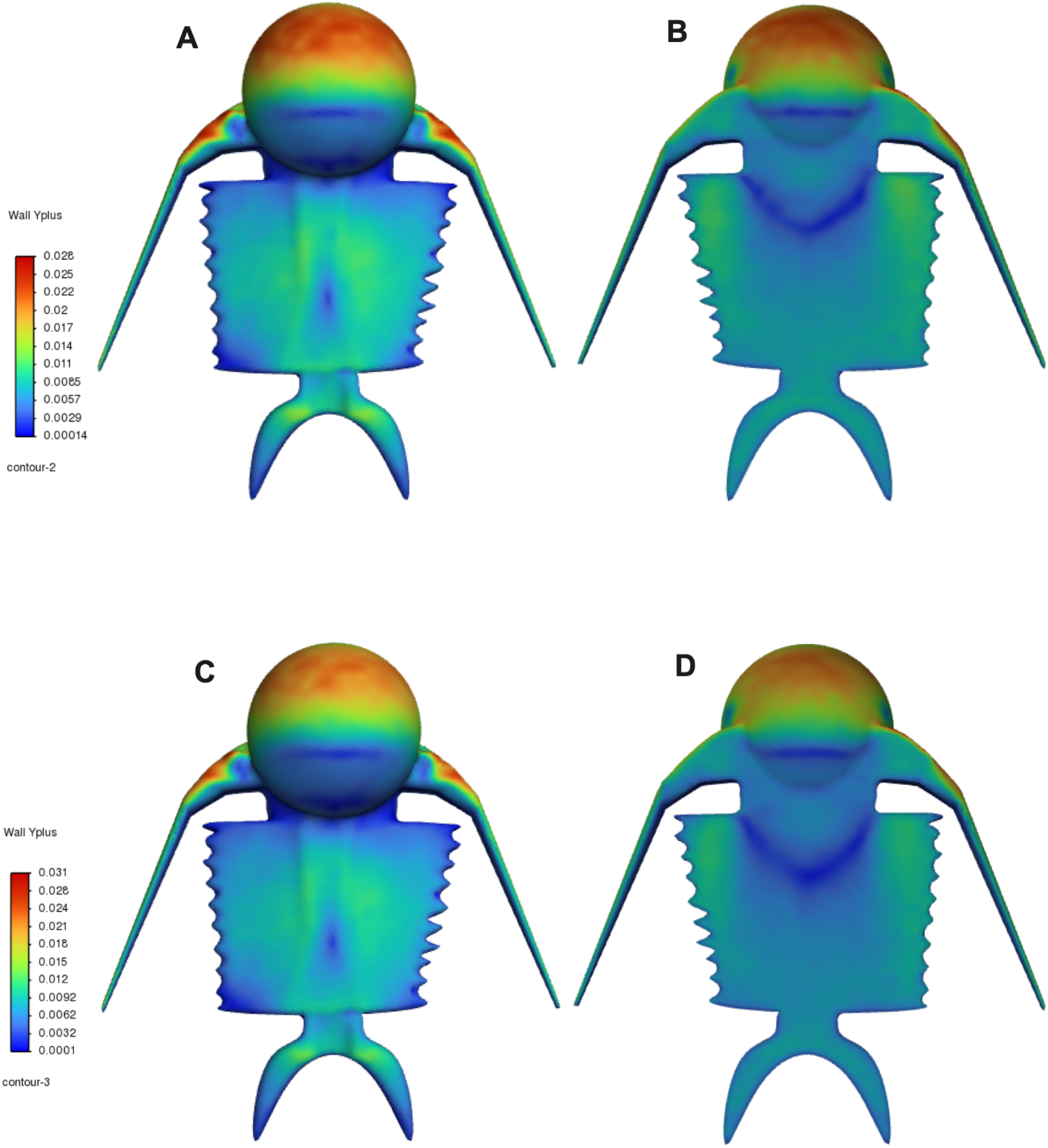
Wall Y plus coefficient on model of *Deiphon*. **A,** Medium mesh dorsal view. **B,** Medium mesh ventral view. **C,** Fine mesh dorsal view. **D,** Fine mesh ventral view.

**Supplemental Figure 3.**
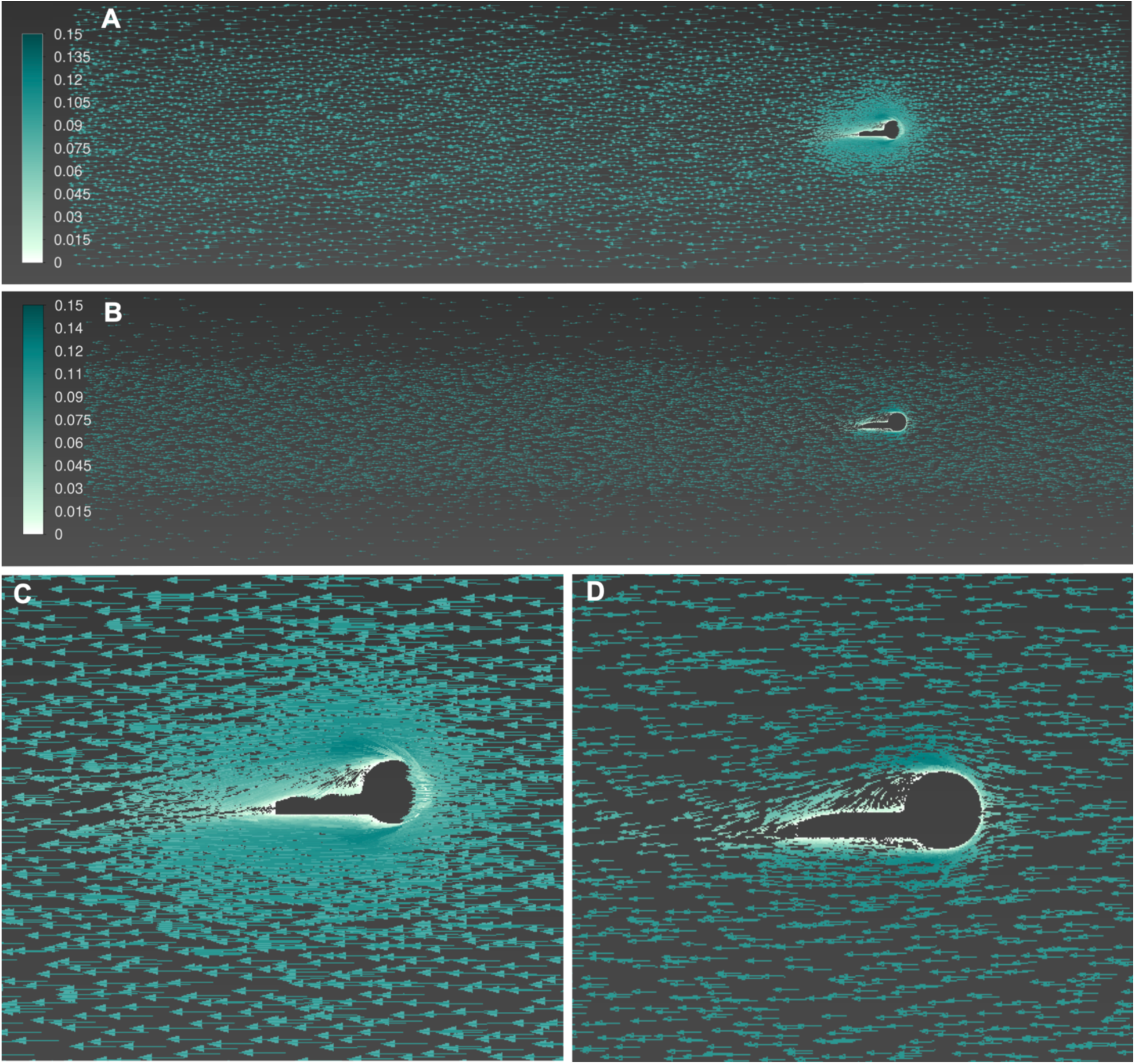
A, Velocity field for CFD simulation using fine mesh. **B,** Velocity field for CFD simulation using course mesh. **C,** Magnification of CFD simulation using fine mesh. **D,** Magnification of CFD simulation using course mesh.

**Supplemental Table 1. Measurements of trilobite exoskeletons and appendages.**

**Supplemental Table 2. Density of modern marine arthropods.**

**Supplemental Table 3. Occurrences of *Deiphon* from PBDB.**

**Supplemental Table 4. Specific gravity of modern arthropods.**

## Notes

### Competing Interest Statement

The authors have declared no competing interest.

